# Abstract deliberation by visuomotor neurons in prefrontal cortex

**DOI:** 10.1101/2022.12.06.519340

**Authors:** Julie A. Charlton, Robbe L. T. Goris

## Abstract

During visually guided behavior, the prefrontal cortex plays a pivotal role in mapping sensory inputs onto appropriate motor plans [1]. When the sensory input is ambiguous, this involves deliberation. It is not known whether the deliberation is implemented as a competition between possible stimulus interpretations [2, 3] or between possible motor plans [4, 5, 6]. Here we study neural population activity in prefrontal cortex of macaque monkeys trained to flexibly report categorical judgments of ambiguous visual stimuli. Our task design allowed for the dissociation of neural predictors of the upcoming categorical choice and the upcoming motor response used to report this choice. We find that the population activity initially represents the formation of a categorical choice before transitioning into the stereotypical representation of the motor plan. We show that stimulus strength and prior expectations both bear on the formation of the categorical choice, but not on the formation of the action plan. These results suggest that prefrontal circuits involved in action selection are also used for the deliberation of abstract propositions divorced from a specific motor plan, thus providing a crucial mechanism for abstract reasoning.

---

Our perceptual interpretation of the environment guides our actions. Actions are constrained by the affordances of particular environmental contexts. In a given context, perceptual interpretations may be stereotypically linked to specific actions. For example, when a driver in congested traffic sees the car ahead slow down, she will lift her foot from the gas pedal. When she sees the car speed up, she will instead press the gas pedal more firmly. Perceptual estimates of car speed are imperfect. Deciding how to act in traffic therefore requires deliberation, especially when the changes in car speed are subtle. Deliberation here refers to the computational process of weighing evidence in favor of different choice options. Under static contextual circumstances, brain regions involved in action selection appear to represent such deliberation processes as a competition among possible action plans [7, 8, 9]. But natural behavior occurs under many different contexts and therefore generally requires a flexible association between perceptual interpretation and motor response. It has been hypothesized that when such flexibility is required, deliberation may consist of a competition among possible interpretations of the sensory environment rather than among possible action plans [10, 11, 12, 13].

Here we test this hypothesis using a task requiring flexible reporting of categorical perceptual decisions. We trained two macaque monkeys (F and J) to judge whether a visual stimulus presented near the central visual field was oriented clockwise or counterclockwise from vertical (Fig. 1a-d). The monkeys communicated their judgment with a saccade to one of two peripheral visual targets. The meaning of each response option was signaled by the target’s orientation (clockwise vs counterclockwise), and was unrelated to its spatial position (one target was placed in the neurons’ estimated motor response field, the other on the opposite side of the fixation mark; see Methods). Because the spatial configuration of the choice targets varied randomly from trial-to-trial, the task requires subjects to flexibly switch between two stimulus-response mapping rules (Fig. 1a). While the animals performed this task, we recorded extracellular responses from neural ensembles in the pre-arcuate gyrus, an area of prefrontal cortex (PFC) involved in the selection of saccadic eye movements [14] that represents visuomotor deliberation [8, 15].

**Figure 1.**
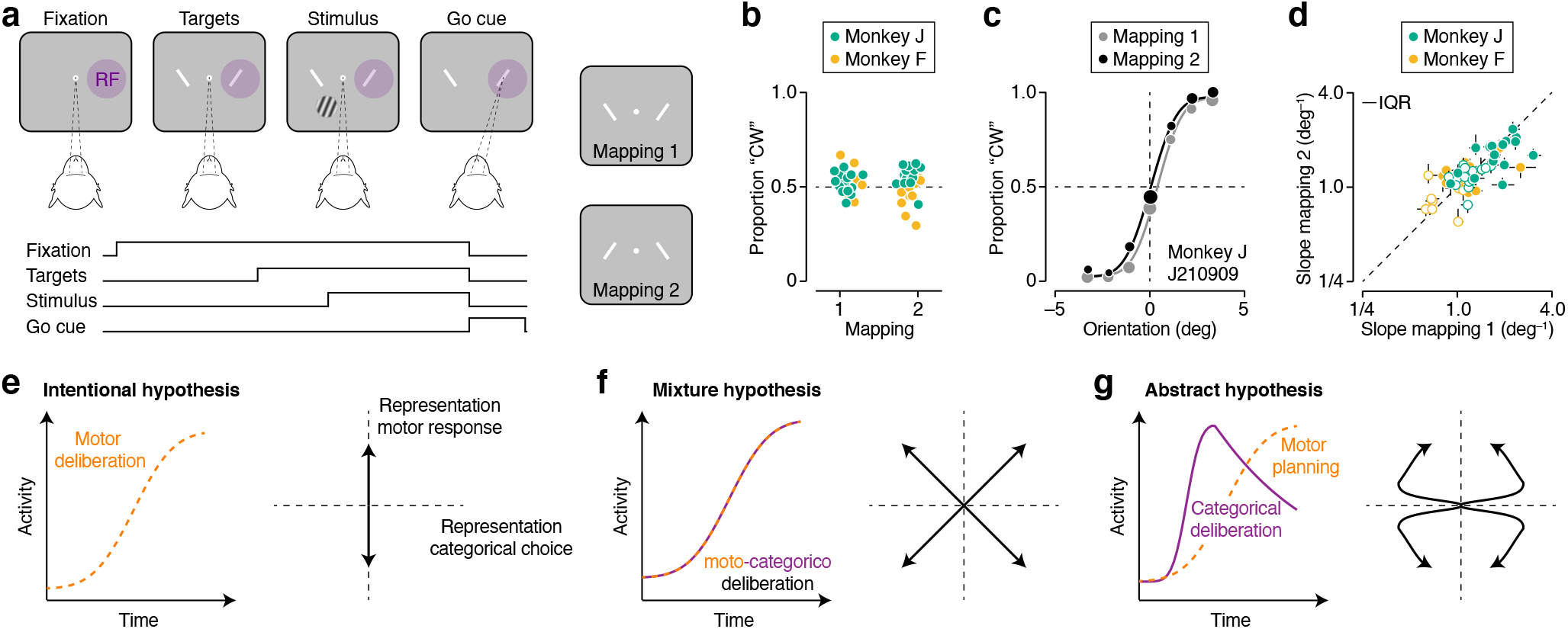
Flexible visual categorization: behavior and computational hypotheses. (**a**) Visual categorization task, task sequence. After the observer fixates for 500 ms, two choice targets appear, followed by the stimulus. The observer judges whether the stimulus is rotated clockwise or counterclockwise relative to vertical and communicates this decision with a saccade towards the matching choice target. Correct decisions are followed by a juice reward. One of the choice targets is placed in the neurons’ presumed motor response field (see Methods). The spatial organization of the choice targets varies randomly from trial-to-trial, giving rise to two stimulus-response mapping rules. (**b**) Proportion of clockwise choices under both mapping rules for both animals. Each symbol summarizes the behavior from a single recording session. (**c**) Psychophysical performance for monkey J in an example recording session. Proportion ‘clockwise’ choices for high contrast stimuli is shown as a function of stimulus orientation under both mapping rules. Symbol size reflects number of trials (total: 1,707 trials). The curves are fits of a behavioral model (see Methods). (**d**) Comparison of orientation sensitivity (i.e., the slope of the psychometric function) under both mapping rules for both monkeys (see Methods). Each symbol summarizes data from a single recording session. Closed symbols: high contrast stimuli, open symbols: low contrast stimuli. Error bars reflect the IQR of the estimate. (**e-g**) Computational hypotheses (left) and associated neural representation motifs (right). There are four possible behavioral outcomes (i.e., either a clockwise or counterclockwise choice, communicated with either a left or rightward saccade), resulting in four motifs per hypothesis.

We found that the activity of many units was not only predictive of the upcoming motor response, but also of the categorical meaning of the choice. Decoding the population activity offered further insight into the evolving decision state of the monkeys. We demonstrate that, following stimulus onset, population activity initially represents the formation of a categorical choice before transitioning into the stereotypical representation of the upcoming motor response. As predicted by theoretical models of decision-making, the formation of the categorical choice reflected a graded representation of evidence, informed by both the current sensory input and stimulus expectations. This was not true of the evolving representation of the motor plan. Our results suggest that prefrontal circuits involved in action selection also support deliberation among abstract propositions.

## Behavior and single unit responses

Both monkeys successfully learned to categorize stimulus orientation under the two mapping rules. Their perceptual choices were evenly distributed among both response alternatives (Fig. 1b), and lawfully depended on stimulus orientation (Fig. 1c). They made few errors in the easiest stimulus conditions (monkey F = ±3.75 deg, median performance = 96.25% correct; monkey J = ±3.3 deg, median performance = 94.38% correct; Extended Data Fig. 1a). The spatial location of the choice targets varied across recording sessions, impacting the animals’ orientation sensitivity. It did so in similar fashion under both mapping rules (median difference in orientation sensitivity: Monkey J = 4.4%, *P* = 0.45; Monkey F = 4.7%, *P* = 0.38; Wilcoxon signed-rank test; Fig. 1d). This pattern was also evident in the animals’ response times (Extended Data Fig. 1b). Together, these results suggest that, within each session, the quality and duration of the decision process did not meaningfully vary across the two mapping rules.

What is the nature of the decision process that underlies this flexible behavior? One viable strategy would be to evaluate which saccadic eye movement is more likely to be correct (the “intentional” hypothesis; Extended Data Fig. 2). In principle, this strategy can be instantiated by oculomotor neural circuits. Alternatively, the deliberation may concern which categorical choice option is most likely to be correct (the “abstract” hypothesis; Extended Data Fig. 2). However, it is not clear which neural circuits would instantiate this computation. Finally, the deliberation process might involve joint consideration of the stimulus category and the corresponding motor plan (the “mixture” hypothesis; Extended Data Fig. 2). We designed the task such that each of these strategies produces a qualitatively distinct ‘motif’ of population activity which represents the unfolding visuomotor deliberation process. The motifs are defined by the joint evolution of activity related to the upcoming categorical choice and the upcoming saccade direction (Fig. 1e–g). We thus set out to characterize the dynamic structure of population activity in PFC while the animals generated this behavior.

Consider the activity of four simultaneously recorded units. We targeted neurons whose motor response field was likely to overlap with one of the choice target locations (see Methods). Grouping trials by saccade direction confirmed that the activity of many units was predictive of the upcoming motor response (Fig. 2a, top, dark vs light orange). Grouping the same trials instead by saccade meaning revealed that the activity of many units was also predictive of the categorical choice (Fig. 2a, top, dark vs light purple). The temporal evolution of choice-related activity differed across units, complicating a functional interpretation (Fig. 2a, bottom). But note that in the majority of cases, categorical selectivity peaked before the go cue (monkey F: 83 of 126 units; monkey J: 243 of 363 units), while motor selectivity peaked after the go cue (monkey F: 79 of 126 units; monkey J: 239 of 363 units; Fig. 2b). This pattern suggests that these predictive signals may be separated in time. The same units tended to exhibit both types of choice selectivity. Specifically, the larger the peak motor selectivity was, the larger the peak categorical selectivity tended to be (Fig. 2c; Spearman rank correlation: Monkey J = 0.55, *P* < 0.001; Monkey F = 0.36, *P* < 0.001). However, there was no obvious relationship between the units’ preferred saccade direction and their preferred stimulus category (Extended Data Fig. 3). Such mixed selectivity is thought to offer significant computational advantage over specialized responses for implementing flexible input-output mappings as required for our task [16, 17, 18].

**Figure 2.**
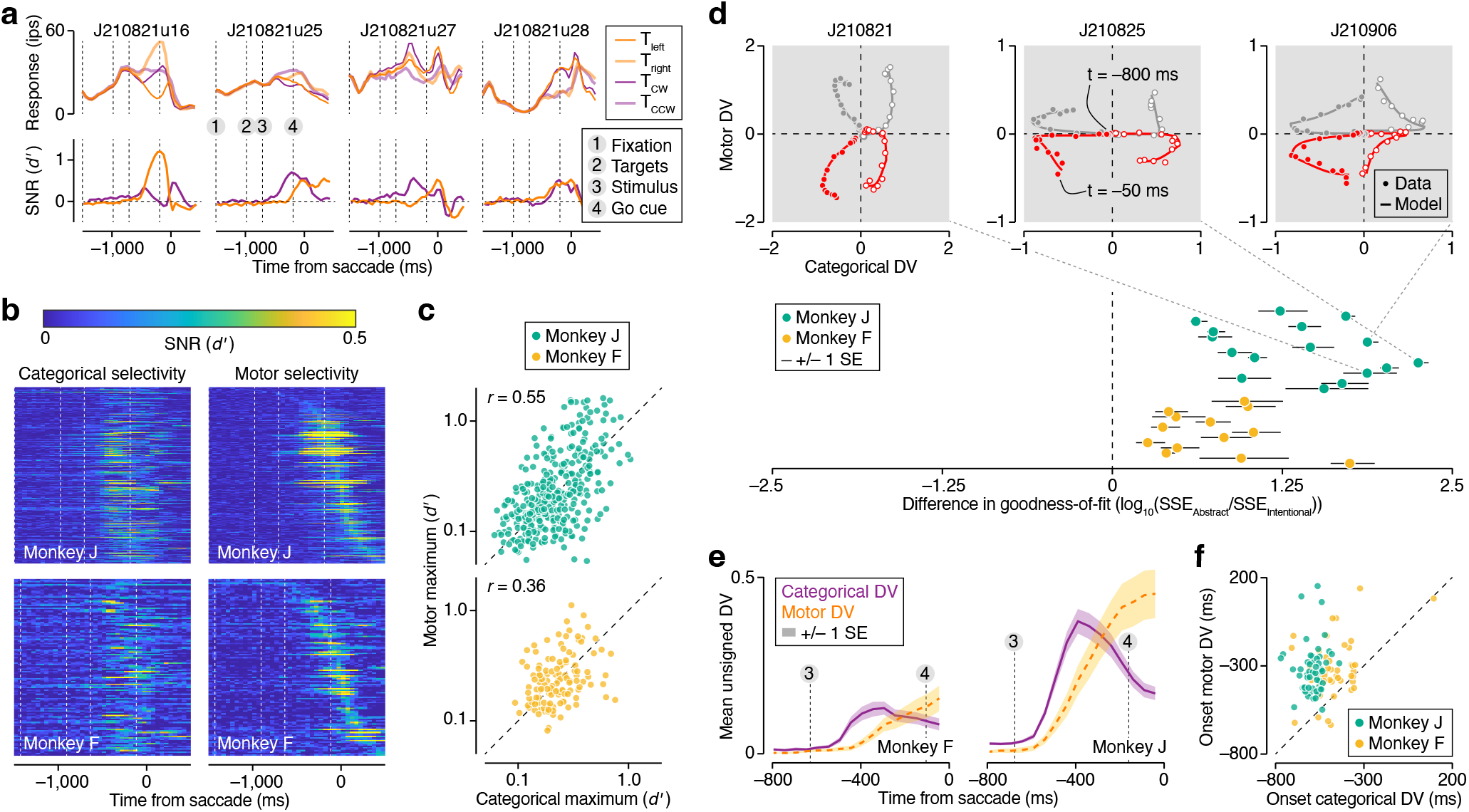
Dynamics of neural activity in PFC during flexible visual deliberation. (**a**) Temporal evolution of firing rate (top) and response selectivity (bottom) of four jointly recorded units (ensemble size: 29 units). Spikes were counted using 50 ms wide counting windows and averaged across trials that either shared the same saccade direction (dark vs light orange), or the same categorical meaning (dark vs light purple). Vertical lines indicate the average time of critical task events. (**b**) Temporal evolution of response selectivity for the categorical choice (left) and the saccade direction (right) of all units recorded from Monkey J (top) and Monkey F (bottom). In all displays, units are ranked according to the timing of their maximal motor selectivity. Vertical lines indicate the average time of critical task events. (**c**) Maximal response selectivity for saccade direction plotted against maximal selectivity for the categorical choice on logarithmic axes. *r* = Spearman correlation. *N* = 363 units for monkey J, and 126 units for monkey F (**d**) Top: Example DV trajectories during a 750 ms epoch preceding saccade initiation for three recording sessions. Symbols represent cross-validated data-based estimates, lines the fit of a descriptive model instantiating the abstract hypothesis (see Methods). Bottom: comparison of goodness-of-fit of two descriptive models instantiating the abstract and intentional hypothesis. Error bars illustrate ±one standard error of the mean, computed across each recording session’s four trajectories. N =16 recording sessions for monkey J, and 13 sessions for monkey F (**e**) Average observed unsigned DV trajectories. Each recording session contributes two unsigned trajectories to this plot. Error bands illustrate ±one standard error of the mean. Vertical lines indicate the average time of critical task events. (**f**) Onset of the motor DV plotted against onset of the categorical DV for all trajectories (see Methods).

## Dynamic population representation motifs

To obtain a perspective on neural population activity during flexible visual categorization, we decoded a time-varying decision variable (DV) from jointly recorded responses (see Methods). This decoded DV indicates how well the subject’s upcoming choice can be predicted from a 50 ms bin of neural ensemble activity [19]. Each behavioral choice is summarized by two independent binary variables: the chosen saccade direction and the corresponding categorical meaning. Likewise, the DV is composed of two independent dimensions. Its temporal structure defines the population representation motif and may thus disambiguate the nature of the decision process (Fig. 1e-g).

Consider the DV trajectories of three example ensembles. To a first approximation, an initial excursion along the categorical dimension is followed by an excursion in the motor dimension (Fig. 2d, top, symbols). Quantitatively, these trajectories are well captured by a model that describes an abstract decision strategy (Fig. 2d, top, curves). In contrast, a model commensurate with an intentional decision strategy provides a poorer fit to the same data as it cannot capture temporal structure in the categorical dimension (Extended Data Fig. 4). This pattern held true for each recorded ensemble (Fig. 2d, bottom; see Methods). To further disambiguate between the abstract and mixture hypotheses, we studied the temporal relationship between the two DV dimensions. Key to the mixture hypothesis is the simultaneous evolution of decision-related activity in both dimensions (Fig. 1f). However, the categorical DV systematically preceded the motor DV. This can be seen in the average unsigned observed DV trajectories, obtained by inverting the trajectories associated with “counter-clockwise” and “left” choices and grouping these with the “clockwise” and “right” trajectories, respectively. In both monkeys, the average unsigned categorical DV begins rising within 150 ms following stimulus onset, well before the average unsigned motor DV begins to rise (Fig. 2e). To investigate whether this pattern was also evident at the level of individual DV trajectories, we fit an unconstrained version of the descriptive model to the data (see Methods). The resulting fits closely resembled the observed data (Extended Data Fig. 5a), allowing us to estimate the onset time of each DV’s rise in a systematic manner (see Methods). In the overwhelming majority of individual model-predicted trajectories, the categorical DV began rising well before the motor DV (Fig. 2f). Restricting these analyses of the DV trajectories to the fully ambiguous stimulus condition (stimulus orientation = 0 deg) yielded similar results, suggesting that these patterns of neural activity are intimately related to the unfolding decision process, rather than to underlying physical stimulus differences as such (Fig. 3).

**Figure 3.**
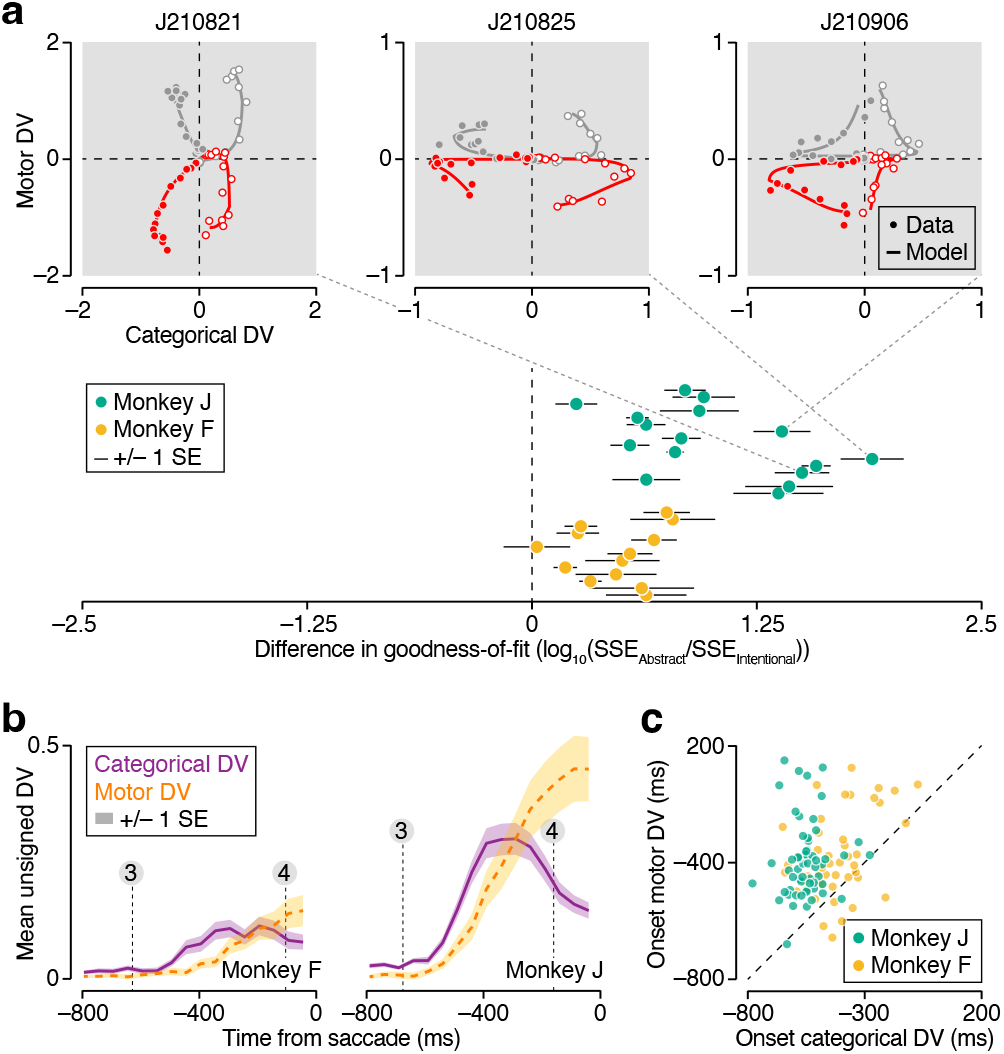
Dynamics of neural activity in PFC during deliberation of zero-signal stimulus. (**a**) Top: Example DV trajectories during a 750 ms epoch preceding saccade initiation for three recording sessions. Only trials that involved the zero-signal stimulus (stimulus orientation = 0 deg) were included in this analysis. Bottom: comparison of goodness-of-fit of two descriptive models instantiating the abstract and intentional hypothesis. Same plotting conventions as Fig. 2d. (**b**) Average observed unsigned DV trajectories (zero-signal trials only). Same plotting conventions as Fig. 2e. (**c**) Onset of the motor DV plotted against onset of the categorical DV for all trajectories (zero-signal trials only).

## Neural signatures of deliberation

We have shown that the temporal structure of population activity in PFC is incompatible with the hypothesis that intentional deliberation underlies the monkeys’ flexible behavior. It is also incompatible with a task-specific variant of this hypothesis (a spatial match-to-sample strategy, see Extended Data Fig. 6), and offers little support for the mixture hypothesis. Instead, our analysis favors the hypothesis that abstract deliberation underlies the monkeys’ flexible behavior. If this interpretation is correct, then the categorical DV ought to exhibit key signatures of deliberation. Moreover, these signatures should not be present in the motor DV. This prediction is unique to the abstract hypothesis (Fig. 1e-g), and thus offers a strong test of our proposed interpretation.

The simplest theoretical models of decision-making hold that subjects solve binary decision-making tasks by comparing the evidence that favors one response alternative over the other with a fixed criterion [20]. Due to noise, repeated presentations of the same stimulus elicit different evidence estimates and may therefore result in different decision outcomes (Fig. 4a, left). When averaged across many trials, this deliberation process gives rise to a graded representation of relative evidence that varies with stimulus strength and differs for correct and incorrect decisions (Fig. 4a, right). For this reason, evidence estimates are thought to not only inform decision outcome, but also determine a subject’s commitment to an evolving decision [8, 9] and factor into their confidence in a decision [21, 22]. If the neural populations we recorded from are involved in the deliberation process, their activity should thus reflect a graded representation of evidence. The issue at stake is whether this representation manifests in the motor DV, the categorical DV, or both.

**Figure 4.**
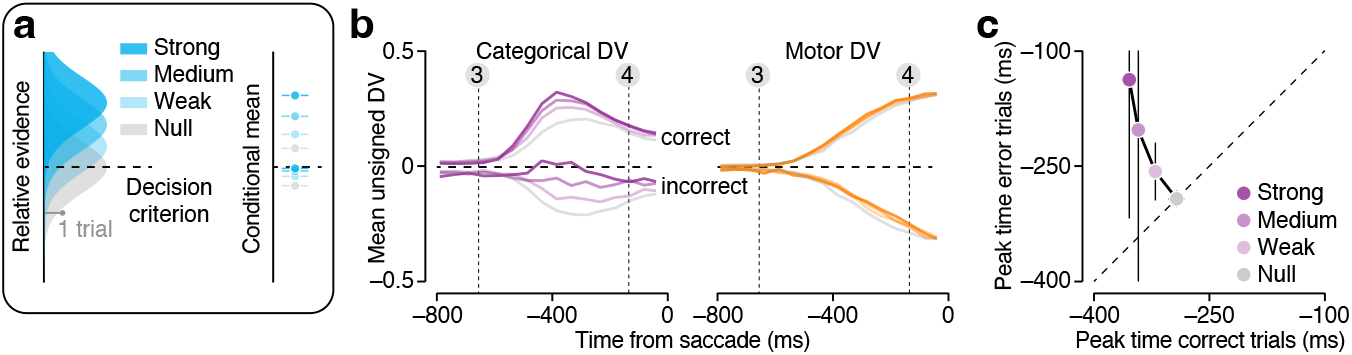
Effects of stimulus strength and choice accuracy on the DV. (**a**) Binary decisions are often modeled as arising from a process whereby relative evidence (i.e., the accumulated evidence that favors one response alternative over the other) is compared to a fixed criterion. (Left) Repeated presentations of the same condition elicit variable amounts of evidence, giving rise to an evidence distribution across many trials (four examples shown). Stronger stimuli result in a smaller overlap of the evidence distribution with the criterion, thus yielding more correct decisions. (Right) The average relative evidence split by choice accuracy (top: correct trials; bottom: incorrect trials) and stimulus strength. (**b**) Average unsigned observed DV trajectories split by choice accuracy (top: correct trials; bottom: incorrect trials) and stimulus strength (i.e., orientation, see Methods). Data of all recording sessions were pooled. Left: Categorical DV. Right: Motor DV. Vertical lines indicate the average time of critical task events as in Figure 2a. (**c**) Comparison of peak time of the average categorical DV trajectories for incorrect and correct trials. Error bars illustrate ±one standard error of the estimate (see Methods).

Consider the temporal evolution of the average unsigned DVs, split by stimulus strength and choice accuracy (Fig. 4b). Dividing trials across this many conditions dilutes the statistical power of the analysis. To compensate for this, we pooled data of both monkeys (see Methods). As can be seen, approximately 150 ms after stimulus onset, the sign and amplitude of the categorical DV begin to match the theoretical prediction of evidence representation. Specifically, the categorical DV achieves more extreme values for correct decisions based on stronger stimuli but exhibits the opposite order for incorrect decisions (Fig. 4b, left). This pattern becomes increasingly prominent over the next 200 ms. The categorical DV trajectories appear to reach their most extreme value more quickly for correct than for incorrect decisions (Fig. 4b, left), consistent with dynamical models of decision-making in which evidence is integrated over time until it reaches a bound [23, 24]. This visual impression was validated by a quantitative analysis (Fig. 4c, Extended Data Fig. 5b; see Methods). In contrast, the amplitude and timing of the motor DV do not appear to reflect the strength of the evidence supporting the choice that informed the upcoming saccade (Fig. 4b, right). The stereotypical nature of the motor DV suggests that it represents a “pure” motor plan.

## Impact of statistical regularities in the environment

Perceptual decisions are not only determined by the present sensory input. They are also shaped by expectations that reflect previously experienced statistical regularities in the environment [25, 26]. Knowledge of such regularities (“prior knowledge”) provides evidence that bears on challenging visual categorization problems. In theory, it can therefore be leveraged to improve the quality of uncertain decisions. Ample empirical evidence demonstrates that humans and other animals heavily exploit prior knowledge for perception [26, 27], action [28, 29], and cognition [30, 31].

We wondered how prior knowledge impacts PFC population representations during flexible visual categorization. To investigate this, we designed the task such that blocks of trials in which clockwise stimuli were over-represented alternated with blocks in which counterclockwise stimuli were over-represented (see Methods). We additionally varied stimulus contrast. The current latent state of each trial was cued to the monkey through the shape of the fixation mark (see Methods). When the stimulus contrast was high, perceptual orientation estimates were more certain, and the impact of the prior on the choice behavior was often small (Fig. 5a, top). When the stimulus contrast was low, perceptual orientation estimates were less certain, as evidenced by the shallowing of the psychometric function (Fig. 5a, bottom; median reduction in orientation sensitivity: Monkey J = 46.4%, *P* < 0.001; Monkey F = 40.7%, *P* < 0.001; Wilcoxon signed-rank test). As a consequence, the impact of the prior on the decision grew, giving rise to increased separation between the prior-specific psychometric functions, hereafter termed “decision bias” (Fig. 5a, top vs bottom; median increase in decision bias: Monkey J = 63.3%, *P* = 0.0013; Monkey F = 68.6%, *P* = 0.04). In general, both monkeys tended to make more biased decisions under task conditions associated with lower orientation sensitivity (Fig. 5b; Spearman rank correlation: Monkey J = −0.42, *P* = 0.017; Monkey F = −0.60, *P* = 0.0015). This trend naturally arises when subjects use the available evidence in a statistically optimal fashion [32, 33].

**Figure 5.**
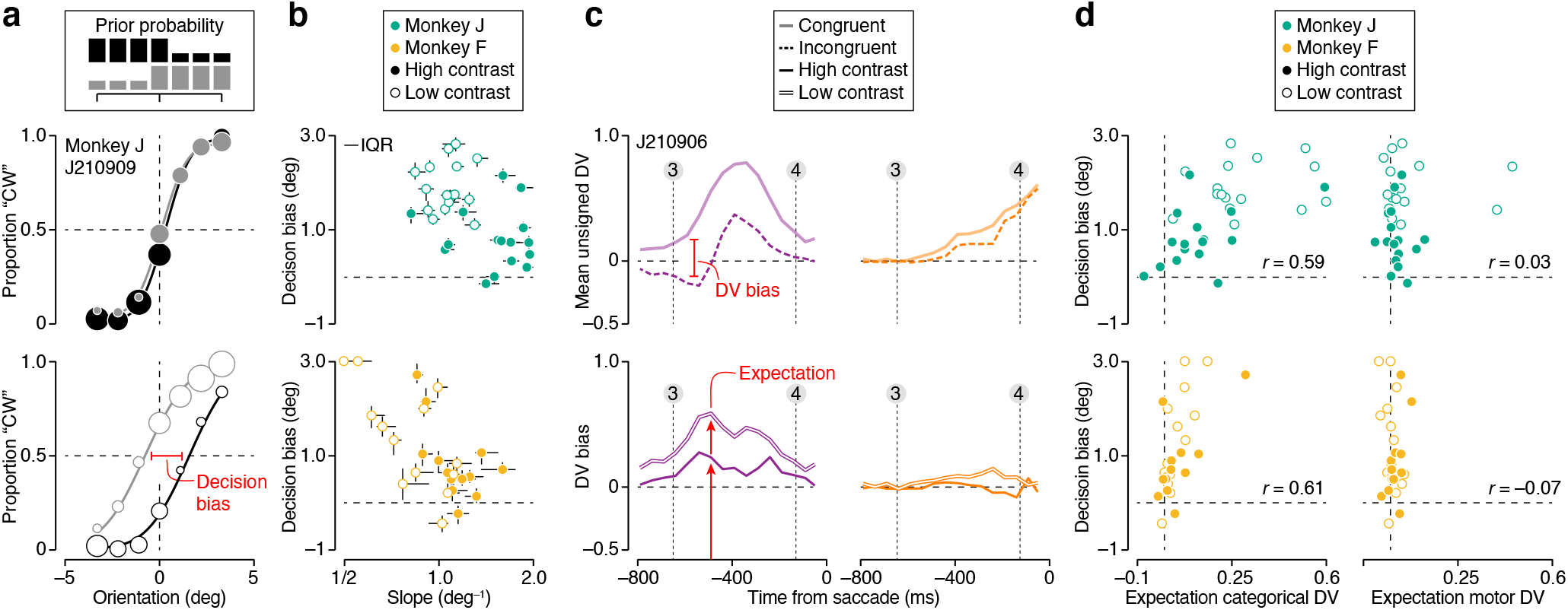
Effects of prior stimulus expectation on the DV. (**a**) Psychophysical performance for monkey J in an example recording session. Proportion clockwise choices is plotted as a function of stimulus orientation under both stimulus priors (black vs grey), split by stimulus contrast (top: high contrast trials, bottom: low contrast trials). Symbol size reflects number of trials (total: 1,707 high contrast trials and 1,875 low contrast trials). The curves are fits of a behavioral model (see Methods). (**b**) Decision bias plotted as a function of orientation sensitivity for both monkeys (top: Monkey J, bottom: Monkey F). Each symbol summarizes data from a single recording session. Closed symbols: high contrast stimuli, open symbols: low contrast stimuli. Error bars reflect the IQR of the estimate. (**c**) Top: Average unsigned DV trajectories split by choice congruency for an example recording session. Only low contrast trials are included. Bottom: DV bias in the example dataset for high and low contrast trials. The categorical DV is shown on the left, the motor DV on the right. (**d**) Decision bias plotted as a function of stimulus expectation for both monkeys. Same plotting conventions as in panel b.

To isolate the effects of the monkeys’ prior knowledge on the neural representation, we compared DV trajectories of trials that resulted in the same categorical choice but that were either congruent or incongruent with the prior expectation (see Methods). As can be seen from an example recording session, congruent and incongruent categorical DV trajectories could differ substantially (Fig. 5c, top left). This difference, which we term DV bias, was often present before stimulus onset and was more prominent during blocks of low-contrast trials (Fig 5c, bottom left). This suggests that it may provide a neural measure of the impact of prior expectations on ensuing perceptual decisions. To test this idea, we calculated the DV bias around the time when the categorical DV first begins to reflect stimulus information (i.e., 500 ms before saccade initiation, Fig. 5c, bottom left, red arrows). For every recording session, we thus obtained two neural measures of “expectation”, one for high contrast trials, and one for low contrast trials. For both monkeys, expectation calculated from the categorical DV predicted the behaviorally measured decision bias (Fig. 5d, left, Spearman rank correlation: Monkey J = 0.59, *P* < 0.001; Monkey F = 0.61, *P* = 0.0011). For the motor DV, this was not the case (Fig. 5d, right, Monkey J = 0.025, *P* = 0.89; Monkey F =–0.067, *P* = 0.74). Calculating neural expectation from slightly earlier or later moments in time yielded similar results (Extended data Fig. 7). These results further corroborate the hypothesis that deliberation occurred in an abstract stimulus representation space. They also imply that during categorical deliberation, PFC activity is not only shaped by input from visual cortex, but also by signals representing prior knowledge retrieved from memory.

## Discussion

In this study, we have investigated neural population activity in PFC during flexible visual categorization. We sought to probe the nature of the decision process that underlies the flexible relationship between perception and action demanded by many of the real-world problems we face. We suggest that behavioral reports arise from a decision process in which evaluating the sensory environment and planning to act on that interpretation are supported by the same populations of neurons, but unfold in separate representational spaces and different moments in time. This view explains three distinct observations. First, during sensory stimulation, an initial population representation of the upcoming categorical choice precedes an orthogonal representation of the motor action used to communicate that choice (Fig. 2–3). Second, neural activity patterns predictive of the upcoming categorical choice reflect a graded representation of evidence, while activity patterns predictive of the upcoming motor response do not (Fig. 4). And third, prior stimulus expectations shape the formation of the categorical choice but not the formation of the action plan (Fig. 5).

Our investigation is the first to offer unequivocal evidence that circuits involved in action selection can also reflect deliberation among abstract propositions in a representational space that is uncoupled from specific motor plans [13]. Previous attempts to determine whether action-planning circuits in the macaque brain also support abstract deliberation were inconclusive for a variety of reasons. Some studies used a temporal match-to-sample task [34, 2, 35]. In these tasks, the decision variable consists of a comparison of two stimulus representations. As a consequence, such tasks allow for the identification of abstract perceptual representations [34, 2, 35], but not for the identification of neural deliberation signals. Some other studies used a task design similar to ours, but found that animals appeared to adopt an intentional strategy and that neural activity did not reflect categorical choice formation [36, 37]. Finally, in most previous studies, neural signals were recorded from one unit at a time and could thus not reveal the structure of population activity [12, 38]. As such, our experimental paradigm opens new possibilities to further investigate the neural basis of abstract perceptual reasoning.

Decision-related activity has been found in many different brain areas [39]. It has been challenging to ascribe a unique role to each of these areas. This requires experimental paradigms that are simple enough to invite well-controlled, reliable behavior, but complex enough to engage higher cognitive mechanisms. Our paradigm revealed dissociable signatures of stimulus strength, perceptual uncertainty, prior knowledge, and action plans within a single area. Our approach therefore holds promise to disambiguate the functional roles of brain areas within the decision-making network, and more generally, to characterize the cascade of neural operations that collectively transform sensory inputs into perceptual interpretations and context-appropriate action plans.

## METHODS

### 0.1 Subjects

Our experiments were performed on two adult male macaque monkeys (*Macaca mulatta*, ages 8-9 years old over the course of the experiments). The animals were trained to perform a memory-guided saccade task and an orientation discrimination task with saccadic eye movements as operant responses. They had not previously participated in research studies. All training, surgery, and recording procedures conformed to the National Institute of Health Guide for the Care and Use of Laboratory Animals and were approved by The University of Texas at Austin Institutional Animal Care and Use Committee. Under general anesthesia, both animals were implanted with three custom-designed titanium head posts and a titanium recording chamber [40].

### 0.2 Apparatus

The subjects were seated in a custom-designed primate chair in front of a CRT monitor (Sony Trinitron, model GDM-FW900), with their heads restrained using three surgical implants. Stimuli were shown on the CRT monitor which was positioned approximately 64 cm away from the monkeys’ heads. Eye position was continuously tracked with an infrared eye tracking system at 1 kHz (Eyelink 1000, SR Research, Canada). Stimuli were generated using the Psychophysics Toolbox [41] in MATLAB (MathWorks). Neural activity was recorded using the Plexon OmniPlex System (Plexon). Precise temporal registration of task events and neural activity was obtained through a Datapixx system (Vpixx). All of these systems were integrated using the PLDAPS software package [42].

### 0.3 Memory-guided saccade task

We used a variation of the classical memory-guided saccade task [43] to identify recording sites where neurons exhibited neural activity indicative of an upcoming eye movement. Each trial began when the subject fixated a small white square at the center of the screen. After 100 ms, a small response target briefly appeared in one of 24 possible locations (3 radii x 8 directions). The subject needed to keep this location in memory while maintaining fixation for 500 ms. After this delay period, the fixation mark disappeared and the subject needed to make a saccade to the remembered location. Correct choices were followed by a juice reward. Each location was presented multiple times per recording session.

### 0.4 Estimating response field locations

During the memory-guided saccade task, extracellular recordings were made with dura-penetrating glass-coated tungsten microelectrodes (Alpha Omega), advanced mechanically into the brain. We made recordings from multiple sites in the pre-arcuate gyrus. After data collection was completed, we studied spiking activity in a 100 ms window preceding saccade initiation. We compared the strength of the response preceding an eye movement to the neuron’s apparent preferred spatial location with the responses preceding eye movements to all other locations. We deemed a neuron to have a well-defined motor response field if this difference fell outside the expected difference distribution predicted by a null-model that assumes Poisson spiking statistics. Following identification of a suitable recording site, we conducted several additional orientation discrimination training sessions with one choice target placed within the estimated response field location and one on the opposite site of the fixation mark. Once psychophysical performance reached a high level, physiological data collection begun.

### 0.5 Orientation discrimination task

The orientation-discrimination task is a variant of classical visual categorization tasks in which the subject uses a saccadic eye movement as operant response [44, 45, 46]. We used a flexible version of this task in which the stimulus-response mapping rule varied from trial to trial. Each trial began when the subject fixated a small white square at the center of the screen (0.6 degrees in diameter). Upon fixation, the square was replaced by either a triangular or a circular fixation mark, indicating the latent prior context of the trial. The experiment involved two distinct prior contexts, associated with differently skewed distributions of stimulus orientation (see inset of Fig. 5a). Blocks of both priors alternated randomly (80 trials per block). 500 ms + 0-65 ms after the onset of the fixation mark, two choice targets appeared, one on each side of the fixation mark. One choice target was placed within the presumed motor response field, the other on the opposite side of the fixation mark. The choice targets were white lines (2.5 deg x 0.5 deg), rotated –22.5 deg and 22.5 deg from vertical. 250 ms + 0-65 ms later, a circularly vignetted drifting grating appeared in the near periphery (eccentricity: 1.12 degrees). The grating measured 2.7 degrees in diameter, had a spatial frequency of 1 cycle/deg, and a temporal frequency of 1 cycle/s. The stimulus remained on for 500 ms + 0-65 ms. Subjects judged the orientation of the stimulus relative to vertical. The stimulus then disappeared along with the fixation mark and subjects reported their decision with a saccadic eye movement to the appropriately oriented choice target.

Trials in which the monkey did not saccade to either of the choice targets within 2 s were aborted. Auditory feedback about the accuracy of the monkey’s response was given at the end of each trial. Correct choices were followed by a liquid reward delivered via a solenoid-operated reward system (New Era). Stimulus orientation varied over a small range, tailored to each monkey’s orientation sensitivity (monkey F: −3.75 deg to 3.75 deg, monkey J: −3.3 deg to 3.3 deg). Vertically oriented stimuli received random feedback. Stimuli were presented at either high or low contrast (Michelson contrast: 100% or 4%). Blocks of high and low contrast stimuli alternated randomly (trials per block: monkey F = 100, monkey J = 80). We conducted 13 successful recordings from monkey F and 16 from monkey J (average number of trials per session, monkey J = 3,171; monkey F = 1,593).

### 0.6 Behavioral analysis

We measured observers’ behavioral capability to discriminate stimulus orientation by fitting the relationship between stimulus orientation and probability of a “clockwise” choice with a psychometric function consisting of a lapse rate and a cumulative Gaussian function. Model parameters were optimized by maximizing the likelihood over the observed data, assuming responses arise from a Bernouilli process. Each recording session was analyzed independently. For the analysis documented in Fig. 1d, we fit one psychometric function per mapping rule and contrast level. We defined orientation sensitivity as the inverse of the SD of the cumulative Gaussian. We used a variant of this model to measure observers’ prior-induced behavioral decision bias. For this analysis, we fit one psychometric function per stimulus prior and contrast level (Fig. 5a). Both prior conditions shared the same sensitivity parameter, resulting in two psychometric functions with identical slope. We defined decision bias as the difference between the means of both cumulative Gaussians (i.e., the magnitude of the horizontal displacement of both psychometric functions). Error bars of model-based statistics are based on a 100-fold non-parametric bootstrap of the behavioral data.

### 0.7 Electrophysiological recordings

During the orientation-discrimination task, we recorded extracellular spiking activity from populations of PFC neurons through a chronically implanted recording chamber. Every recording session, we used a microdrive (Thomas recording) to mechanically advance a linear electrode array (Plexon S-probe; 32 contacts) into the brain at an angle approximately perpendicular to the cortical surface. We targeted recording sites that had exhibited well-defined motor response fields in a previously conducted memory-guided saccade task. We positioned the linear arrays so that they roughly spanned the cortical sheet and removed them after each recording session. Continuous neural data were acquired and saved to disk from each channel (sampling rate 30 kHz, Plexon Omniplex System). To extract responses of individual units, we performed offline spike sorting. We first automatically spike-sorted the data with Kilosort [47], followed by manual merging and splitting as needed. Given that the electrode’s position could not be optimized for all contact sites, most of our units likely consist of multi-neuron clusters. All units whose mean firing rate during the task exceeded 3 ips were included in the analysis.

### 0.8 Analysis of single unit responses

We measured the temporal evolution of each unit’s response by expressing spike times relative to the trial-specific moment of saccade initiation and counting spikes within non-overlapping 50 ms windows. Fig. 2a shows example response traces for four units, averaged across different subsets of trials. We computed neuronal selectivity for the upcoming choice behavior by calculating the difference between the choice-conditioned response averages, normalized by the response standard deviation [48]. The sign of this SNR metric depends on the unit’s preferred choice option. To facilitate comparison across the categorical and motor dimension, we signed each unit’s SNR-trace such that the maximal value was positive (see examples in Fig. 2a, all traces are shown in Fig. 2b).

### 0.9 Estimating the time-varying decision variable

For each trial, we obtained moment-to-moment measurements of the decision variable by projecting 50 ms bins of population activity onto a linear decoder optimized to distinguish the activity patterns associated with both choice options (“left” vs “right” choices for the motor DV, and “clockwise” vs “counterclockwise” choices for the categorical DV, respectively). Specifically, we first individually z-scored each unit’s spike counts within every time bin. We then used these z-scored responses to estimate the set of linear weights, **w** = (*w*_1_,…, *w_n_*), that best separate the choice-conditioned z-scored response patterns, assuming a multivariate Gaussian response distribution:

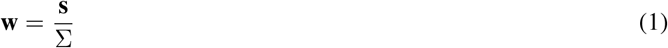

where **s** is the mean difference of the choice-conditioned z-scored responses and Σ is the covariance matrix of the z-scored responses. The decoder weights are calculated from observed trials. To avoid double-dipping, we excluded the trial under consideration from the calculation and solely used all other trials to estimate the weights. This way, we obtained “cross-validated” DV estimates for each time bin:

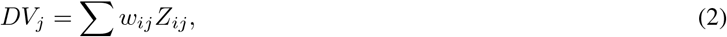

where *W_ij_* and *Z_ij_* are the weight and z-scored response of unit *i* on trial *j* for a given time bin. The symbols in Fig. 2d show DV trajectories from three example recording sessions, averaged across all choice-conditioned trials. The symbols in Fig. 3a show DV trajectories from the same example recording sessions for the zero-signal stimulus. The lines in Fig. 2e and Fig. 3b show unsigned DV trajectories, obtained by inverting the trajectories associated with “counter-clockwise” and “left” choices and grouping these with the “clockwise” and “right” trajectories, respectively. The lines in Fig. 4b show unsigned DV trajectories, split by stimulus strength and choice accuracy, and averaged across all recording sessions of both animals. The lines in the top panel of Fig. 5c show unsigned DV trajectories of an example recording session averaged across choice “congruent” and “incongruent” trials, respectively.

### 0.10 Descriptive models of computational hypotheses

We compared the observed DV trajectories with the theoretical expectations of two computational models of decision-making. We expressed the models’ predictions using a set of equations that describe the average evolution of the choice-conditioned decision variable. Under the intentional model, the categorical DV has no systematic structure while the motor DV evolves according to a cumulative Gaussian function. This model has four free parameters per choice-conditioned trajectory: one captures an initial offset in the motor DV, one specifies the dynamic range of the DV trajectory, one controls the speed of the rise, and one the time point at which half of the rise is completed. Under the abstract model, an initial rise in the categorical DV is followed by a subsequent rise of the motor DV. Following completion of the deliberation process, the categorical DV may decay in strength. We used nine free parameters to describe this pattern. Five of these specify the evolution of the categorical DV, and four that of the motor DV. For both DVs, we used cumulative Gaussians in the same way as we did for the intentional model. For the categorical DV, we additionally used a parameter that controls the amount of decay that follows the peak of the categorical DV (defined as the time at which the cumulative Gaussian reached the 99.38th percentile). We imposed boundaries on the model’s parameters that ensured that the motor DV could not begin to rise before the categorical DV. We fit both descriptive models by minimizing the sum of the square error of the choice-conditioned trajectory under consideration. Example fits of the abstract model are shown in Fig. 2d and Fig. 3a, example fits of the intentional model are shown in Extended Fig. 4.

### 0.11 Estimating onset and peak time of DV trajectories

We conducted an analysis in which we compared the estimated onset time of both DVs (Fig. 2f and Fig. 3c). We obtained estimates of onset time by fitting an unconstrained version of the descriptive model to the data. This model used the same set of equations as the abstract model, but we imposed no boundaries on the model’s parameters that would enforce a temporal order on the DV trajectories. The average fit of this model to the data is shown in Extended Data Fig. 5a. For each DV trajectory, we defined onset time as the time at which the cumulative Gaussian reached the 5th percentile. We also conducted an analysis in which we compared the estimated peak time of the categorical DV for different groups of trials (Fig. 4c). We obtained estimates of peak time by fitting the same unconstrained version of the model to each trajectory shown in Fig. 4b. The fits are shown in Extended Fig. 5b. Under this model, peak time is defined as the time at which the cumulative Gaussian reaches the 99.38th percentile (at this time, the decay begins). We obtained estimates of the standard error by repeating this analysis on 1,000 matching synthetic data-sets, each created by sampling the observed trials with replacement. We then performed the entire analysis sequence on these bootstrapped trials. The error bars in Fig. 4c show the estimate for the observed data ±one standard deviation of the peak time estimates of the synthetic data-sets.

### 0.12 Estimating DV bias

We obtained estimates of DV bias by first calculating the average observed unsigned DV trajectory for congruent and incongruent trials per level of stimulus strength (i.e., rotation magnitude), then taking the difference of these averages per level, and finally averaging across these differences. This estimation procedure ensures that stimulus strength as such does not impact the bias estimate (the fraction of congruent and incongruent choices differs across stimulus strengths).

## Acknowledgements

We thank Z. Boundy-Singer, B. Cumming, A. Hermundstad, N. Priebe, and C. Ziemba for helpful comments on an early version of this manuscript. We are grateful to Q. Jeffs for animal support and C. Badillo for technical support throughout the project. This work was supported by US National Institutes of Health grants T32 EY021462 (J.A.C.), EY032999 (R.L.T.G.), and National Science Foundation CAREER award #2146369 (R.L.T.G.).

## Author contributions

J.A.C. and R.L.T.G. conceived and designed the study. J.A.C. collected the data. J.A.C. and R.L.T.G. analyzed the data. J.A.C. and R.L.T.G. wrote the manuscript.

## Competing Interests

The authors declare no competing interests.

---

**Extended Data Figure 1.**
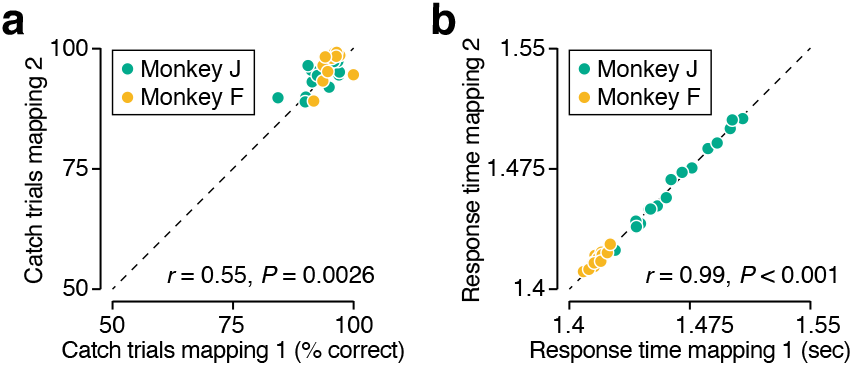
Further comparison of psychophysical performance under both mapping rules. (**a**) Proportion correct judgements for the easiest stimulus conditions (i.e., the two most extreme stimulus orientations). Only high contrast trials were included in the analysis. Each symbol summarizes the behavior from a single recording session. Task performance consistently approached the level expected from a flawless observer without attentional lapses (i.e., 100% correct) and did not differ across both mapping rules (median difference in task performance: 1.6%, *P* =1, Wilcoxon signed-rank test). The positive association across both mapping rules indicates that the fraction of guesses may vary across sessions, but is stable across mapping rules. (**b**) The average response time across all trials completed within a single recording session. Response time is measured relative to the start of the trial. r = Spearman correlation.

**Extended Data Figure 2.**
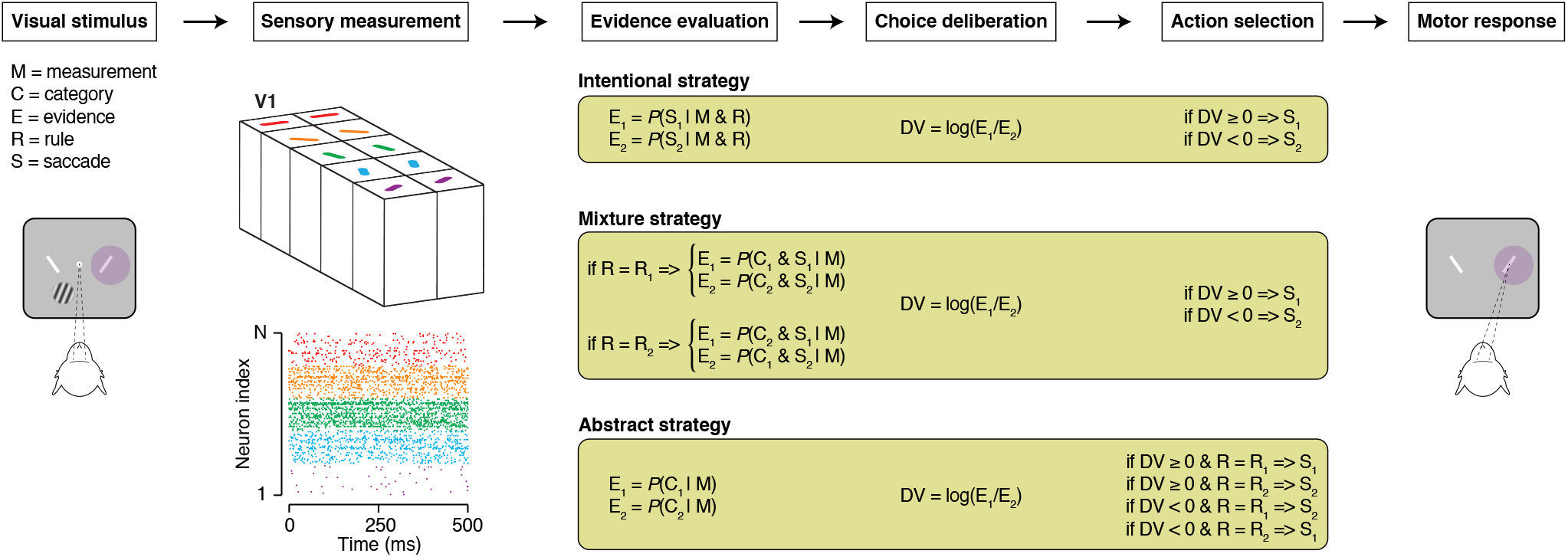
Further comparison of candidate computational strategies. The sensorimotor transformation underlying flexible visual categorization in our task can be broken down into a sequence of conceptually distinct operations (top, boxes). Common to all three candidate strategies is that information about stimulus orientation must be obtained from a sensory measurement (left part of diagram) and that the decision must be communicated with a saccadic eye movement (right part of diagram). The sensory measurement is likely provided by the population activity of a set of visual neurons whose responses selectively depend on stimulus orientation (e.g., by the collective output of a cortical hypercolumn in primary visual cortex). Under an intentional strategy, this activity is evaluated by converting it into evidence in favor of each possible motor response (E_1_ and E_2_, which ideally reflect the likelihood of each response option being correct). This transformation requires taking into account the trial-specific mapping rule. Under an abstract strategy, the sensory activity is evaluated by converting it into evidence in favor of each possible categorical response. This transformation does not require knowledge of the mapping rule. Under a mixture strategy, sensory activity is transformed into evidence favoring one of two possible combinations of categorical choice and associated saccade option. The mapping rule determines the trial-specific pair of combinations. Under all three strategies, choice deliberation involves comparing the evidence in favor of each response option. The logarithm of the likelihood ratio provides a principled metric for this operation. Under the intentional and mixture strategy, the deliberation process directly results in a motor plan. Under the abstract strategy, following deliberation, the mapping rule must be consulted to form the appropriate motor plan.

**Extended Data Figure 3.**
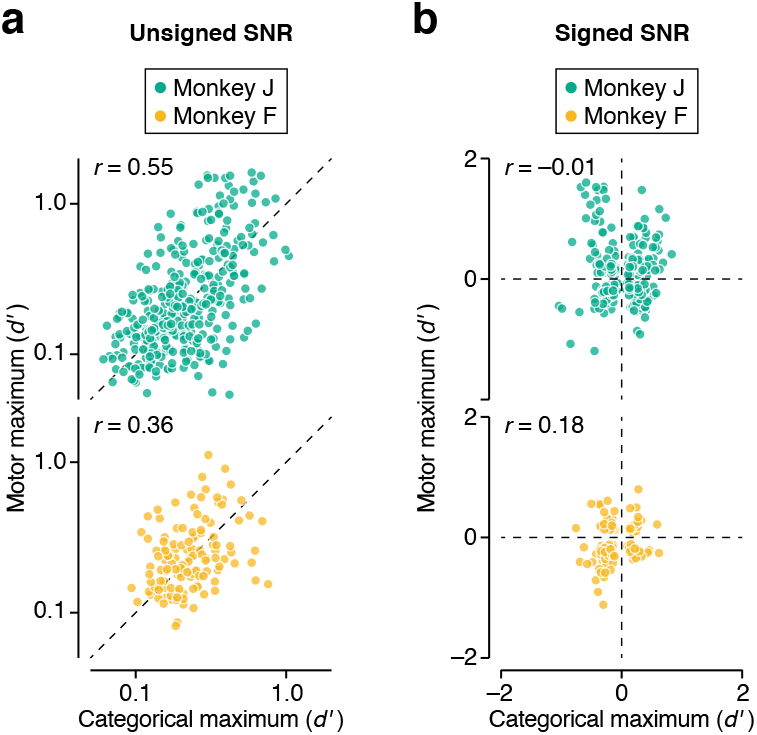
PFC neurons exhibit mixed selectivity for stimulus category and saccade direction. (**a**) Maximal unsigned response selectivity for saccade direction plotted against maximal selectivity for the categorical choice on logarithmic axes (same as Fig 2c of the main paper). (**b**) Most extreme signed response selectivity for saccade direction plotted against maximal selectivity for the categorical choice on linear axes. For both monkeys, every quadrant in the plot is occupied.

**Extended Data Figure 4.**
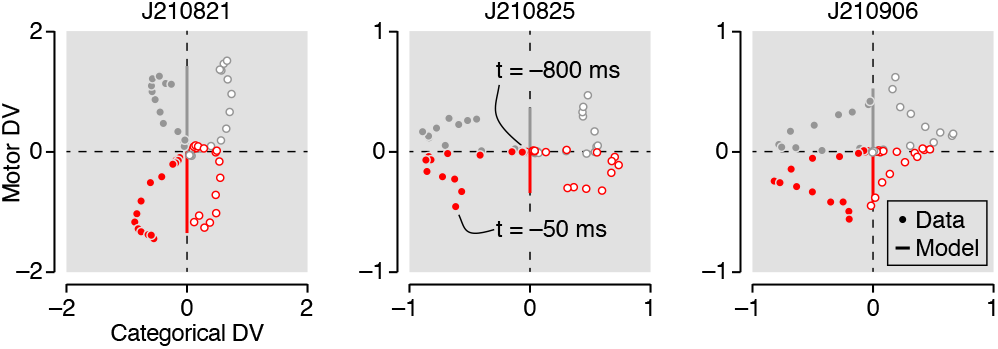
Example DV trajectories during a 750 ms epoch preceding saccade initiation for three recording sessions. Symbols represent cross-validated data-based estimates, solid vertical lines the fit of a descriptive model commensurat¢ with an intentional strategy. This model cannot capture temporal structure in the categorical dimension and hence provides; poor fit to the data (compare with the fit of the abstract model to the same data, shown in Fig. 2d of the paper).

**Extended Data Figure 5.**
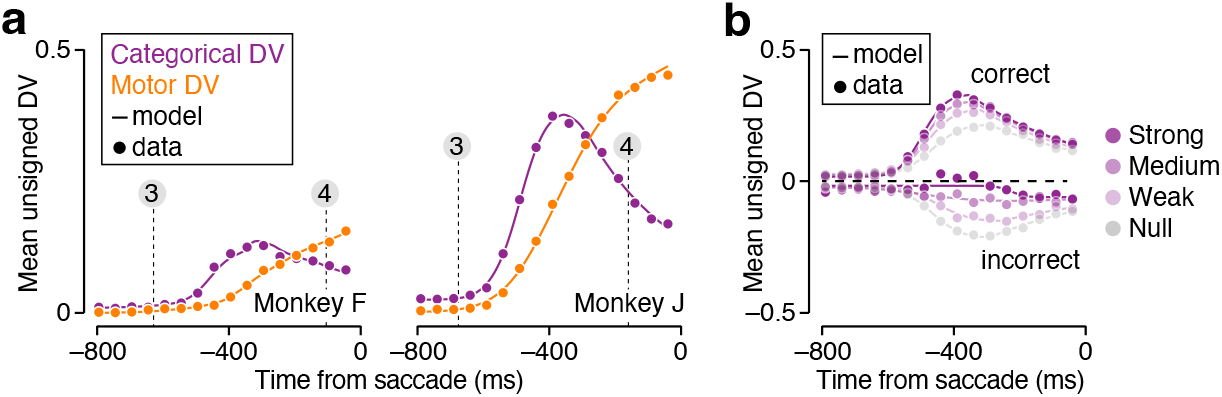
Comparison of model-predicted and observed DV trajectories. (a) Lines show the average unsigned DV trajectories predicted by an unconstrained descriptive model. Each recording session contributes two unsigned trajectories to this plot. Symbols show the average observed values (same data as plotted in Fig 2e in the main paper). Vertical lines indicate the average time of critical task events. These model fits were used to estimate the onset time for each trajectory (shown in Fig. 2f and 3c). (b) Lines show the fit of a descriptive model to the average unsigned DV trajectories split by choice accuracy (top: correct trials; bottom: incorrect trials) and stimulus strength (i.e., orientation). Symbols show the average observed values (same data as plotted in Fig. 4b in the main paper). These model fits were used to estimate the peak time of each trajectory (shown in Fig. 4c).

**Extended Data Figure 6.**
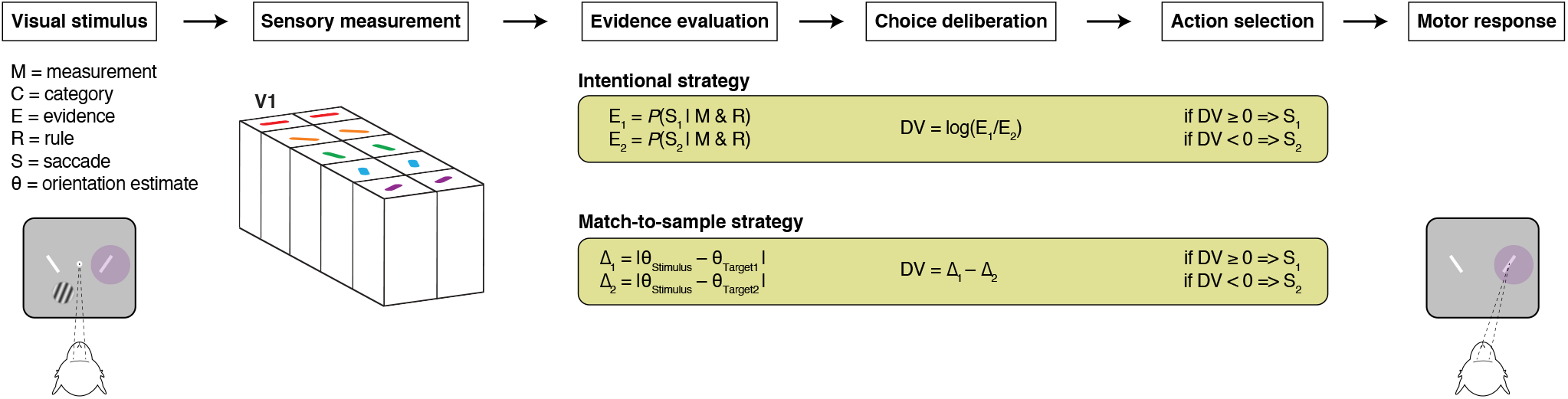
Further comparison of candidate computational strategies. In principle, the subject could solve the task using a spatial match-to-sample strategy. Under this strategy, the perceived stimulus orientation is compared with the orientation of both choice targets, and the most similarly oriented choice target is selected. This strategy is a task-specific variant of an intentional strategy in the sense that the deliberation concerns the question of whether one possible saccade response is favored over the other possible saccade response. Like the intentional hypothesis discussed in the paper, this strategy predicts a data pattern incompatible with our analysis. Specifically, the same stimulus orientation should give rise to oppositely signed DV values under both mapping rules. As documented in the paper, we only see evidence for such a pattern late in the trial, and this pattern does not exhibit neural signatures of deliberation.

**Extended Data Figure 7.**
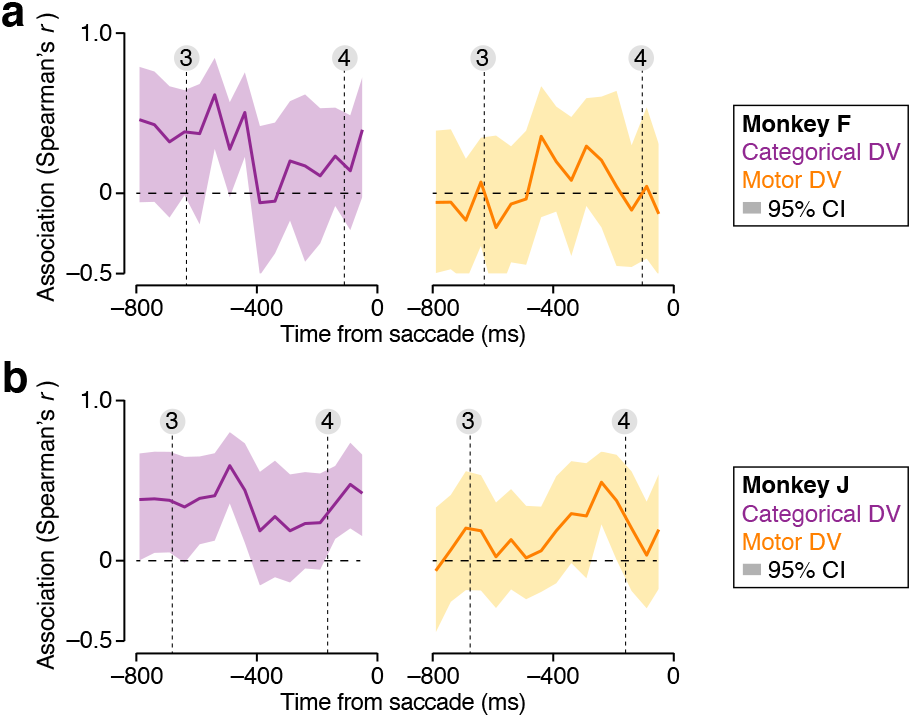
Temporal evolution of the association between a neural measure of expectation and behavioral decision bias. We performed the analysis shown in Fig. 5d of the main paper using a sliding 50 ms wide counting window. For both monkeys, the association between neural expectation calculated from the categorical DV and the behaviorally measured bias was substantial around the time of stimulus onset (indicated by the leftmost dotted line), but decreased as the trial progressed. The association between neural expectation calculated from the motor DV and behavioral bias was minimal around the time of stimulus onset, but gradually increased in strength as the trial progressed. Confidence intervals are based on a 10,000 fold bootstrap test.

## Notes

### Competing Interest Statement

The authors have declared no competing interest.

### Summary of Updates

Error in citation 22 fixed

## References

[1] Earl K Miller and Jonathan D Cohen. An integrative theory of prefrontal cortex function. Annual review of neuroscience, 24(1):167–202, 2001.

[2] David J Freedman and John A Assad. Experience-dependent representation of visual categories in parietal cortex. Nature, 443(7107):85–88, 2006.

[3] David J Freedman and John A Assad. Neuronal mechanisms of visual categorization: an abstract view on decision making. Annual review of neuroscience, 39:129–147, 2016.

[4] Paul Cisek. Cortical mechanisms of action selection: the affordance competition hypothesis. Philosophical Transactions of the Royal Society B: Biological Sciences, 362(1485):1585–1599, 2007.

[5] M Shadlen, R Kiani, T Hanks, and A Churchland. Neurobiology of decision making: An intentional framework in: Engel c, singer w, editors, better than conscious?: decision making, the human mind, and implications for institutions, 2008.

[6] Paul Cisek and John F Kalaska. Neural mechanisms for interacting with a world full of action choices. Annual review of neuroscience, 33:269–298, 2010.

[7] Michael N Shadlen and William T Newsome. Neural basis of a perceptual decision in the parietal cortex (areas lip) of the rhesus monkey. Journal of neurophysiology 86(4):1916–1936, 2001.

[8] Roozbeh Kiani, Christopher J Cueva, John B Reppaŝ, and William T Newsome. Dynamics of neural population responses in prefrontal cortex indicate changes of mind on single trials. Current Biology, 24(13):1542–1547, 2014.

[9] Diogo Peixoto, Jessica R Verhein, Roozbeh Kiani, Jonathan C Kao, Paul Nuyujukian, Chandramouli Chandrasekaran, Julian Brown, Sania Fong, Stephen I Ryu Krishna V Shenoy, et al. Decoding and perturbing decision states in real time. Nature, 591(7851):604–609, 2021.

[10] Joshua I Gold and Michael N Shadlen. The influence of behavioral context on the representation of a perceptual decision in developing oculomotor commands. Journal of Neuroscience, 23(2):632–651, 2003.

[11] Gregory D Horwitz, Aaron P Batista, and William T Newsome. Representation of an abstract perceptual decision in macaque superior colliculus. Journal of neurophysiology, 91(5):2281–2296, 2004.

[12] Sharath Bennur and Joshua I Gold. Distinct representations of a perceptual decision and the associated oculomotor plan in the monkey lateral intraparietal area. Journal of Neuroscience, 31(3):913–921, 2011.

[13] Gouki Okazawa and Roozbeh Kiani. Neural mechanisms that make perceptual decisions flexible. Annual Review of Physiology, 85, 2022.

[14] Jeffrey D Schall. Visuomotor areas of the frontal lobe. In Extrastriate cortex in primates, pages 527–638. Springer, 1997.

[15] Valerio Mante, David Sussillo, Krishna V Shenoy, and William T Newsome. Context-dependent computation by recurrent dynamics in prefrontal cortex. nature, 503(7474):78–84, 2013.

[16] Mattia Rigotti, Omri Barak, Melissa R Warden, Xiao-Jing Wang, Nathaniel D Daw, Earl K Miller, and Stefano Fusi. The importance of mixed selectivity in complex cognitive tasks. Nature, 497(7451):585–590, 2013.

[17] Stefano Fusi, Earl K Miller, and Mattia Rigotti. Why neurons mix: high dimensionality for higher cognition. Current opinion in neurobiology, 37:66–74, 2016.

[18] Alexis Dubreuil, Adrian Valente, Manuel Beiran Francesca Mastrogiuseppe, and Srdjan Ostojic. The role of population structure in computations through neural dynamics. Nature Neuroscience, pages 1–12, 2022.

[19] Yuzhi Chen, Wilson S Geisler, and Eyal Seidemann. Optimal decoding of correlated neural population responses in the primate visual cortex. Nature neuroscience, 9(11):1412–1420, 2006.

[20] W. P. Tanner and J. A. Swets. A decision-making theory of visual detection. Psychological Review, 61:401–409, 1954.

[21] Pascal Mamassian. Visual confidence. Annual Review of Vision Science, 2(1):459–481, 2016.

[22] Zoe M Boundy-Singer, Corey M Ziemba, and Robbe LT Goris. Confidence reflects a noisy decision reliability estimate. Nature Human Behaviour, pages 1–13, 2022.

[23] Richard G Swensson. The elusive tradeoff: Speed vs accuracy in visual discrimination tasks. Perception & Psychophysics, 12(1):16–32, 1972.

[24] Roger Ratcliff and Jeffrey N Rouder. Modeling response times for two-choice decisions. Psychological science, 9(5):347–356, 1998.

[25] H. von Helmholtz. Handbuch der physiologischen Optik, volume III. Leopold Voss, 1867.

[26] Wilson S. Geisler. Visual Perception and the Statistical Properties of Natural Scenes. Annual Review of Psychology, 59(1):167–192, 2008.

[27] Y. Weiss, E. P. Simoncelli, and E. H. Adelson. Motion illusions as optimal percepts. Nature Neuroscience, 5:598–604, 2002.

[28] Konrad P Körding and Daniel M Wolpert. Bayesian decision theory in sensorimotor control. Trends in cognitive sciences, 10(7):319–326, 2006.

[29] Emanuel Todorov. Optimality principles in sensorimotor control. Nature neuroscience, 7(9):907–915, 2004.

[30] Joshua B Tenenbaum, Charles Kemp, Thomas L Griffiths, and Noah D Goodman. How to grow a mind: Statistics, structure, and abstraction. science, 331(6022):1279–1285, 2011.

[31] Thomas L Griffiths, Nick Chater, Charles Kemp, Amy Perfors, and Joshua B Tenenbaum. Probabilistic models of cognition: Exploring representations and inductivebiases. Trends in cognitive sciences, 14(8):357–364, 2010.

[32] Pierre-Simon Laplace. Théorie analytique des probabilités. Courcier, 1812.

[33] Julie A Charlton, Wiktor F Młynarski, Yoon H Bai, Ann M Hermundstad, and Robbe LT Goris. Perceptual decisions exhibit hallmarks of dynamic bayesian inference. bioRxiv, 2022.

[34] David J Freedman, Maximilian Riesenhuber, Tomaso Poggio, and Earl K Miller. Categorical representation of visual stimuli in the primate prefrontal cortex. Science, 291(5502):312–316, 2001.

[35] Chris A Rishel, Gang Huang, and David J Freedman. Independent category and spatial encoding in parietal cortex. Neuron, 77(5):969–979, 2013.

[36] Megan Wang, Christéva Montanède, Chandramouli Chandrasekaran, Diogo Peixoto, Krishna V Shenoy, and John F Kalaska. Macaque dorsal premotor cortex exhibits decision-related activity only when specific stimulus–response associations are known. Nature communications, 10(1):1–16, 2019.

[37] S Shushruth, Ariel Zylberberg, and Michael N Shadlen. Sequential sampling from memory underlies action selection during abstract decision-making. Current Biology, 32(9):1949–1960, 2022.

[38] Yang Zhou and David J Freedman. Posterior parietal cortex plays a causal role in perceptual and categorical decisions. Science, 365(6449):180–185, 2019.

[39] Joshua I Gold, Michael N Shadlen, et al. The neural basis of decision making. Annual review of neuroscience, 30(1):535–574, 2007.

[40] Daniel L Adams, John R Economides, Cristina M Jocson, John M Parker, and Jonathan C Horton. A watertight acrylic-free titanium recording chamber for electrophysiology in behaving monkeys. Journal of neurophysiology, 106(3):1581–1590, 2011.

[41] David H Brainard. The psychophysics toolbox. Spatial vision, 10(4):433–436, 1997.

[42] Kyler M Eastman and Alexander C Huk. Pldaps: a hardware architecture and software toolbox for neurophysiology requiring complex visual stimuli and online behavioral control. Frontiers in neuroinformatics, 6:1, 2012.

[43] Shintaro Funahashi, Charles J Bruce, and Patricia S Goldman-Rakic. Mnemonic coding of visual space in the monkey’s dorsolateral prefrontal cortex. Journal of neurophysiology, 61(2):331–349, 1989.

[44] William T Newsome, Kenneth H Britten, and J Anthony Movshon. Neuronal correlates of a perceptual decision. Nature, 341(6237):52–54, 1989.

[45] Hendrikje Nienborg and Bruce G Cumming. Decision-related activity in sensory neurons may depend on the columnar architecture of cerebral cortex. Journal of Neuroscience, 34(10):3579–3585, 2014.

[46] Robbe LT Goris, Corey M Ziemba, Gabriel M Stine, Eero P Simoncelli, and J Anthony Movshon. Dissociation of choice formation and choice-correlated activity in macaque visual cortex. Journal of Neuroscience, 37(20):5195–5203, 2017.

[47] Marius Pachitariu, Nicholas Steinmetz, Shabnam Kadir, Matteo Carandini, et al. Kilosort: realtime spikesorting for extracellular electrophysiology with hundreds of channels. BioRxiv, page 061481, 2016.

[48] David Marvin Green, John A Swets, et al. Signal detection theory and psychophysics, volume 1. Wiley New York, 1966.

